# Breast cancer through the lens of whole transcriptome spatial imaging

**DOI:** 10.64898/2025.12.19.695622

**Authors:** Claire Williams, Yi Cui, Michael Patrick, Giang Ong, Terence Theisen, Megan Vandenberg, Joseph M. Beechem, Patrick Danaher

## Abstract

Using the world’s first transcriptome-scale spatial imaging of a breast tumor, we assessed what could be learned about one patient’s disease. We cataloged heterogeneity across 3 morphological regions, 9 spatial domains, 37 cell types, and 1692 pathways. Then, employing a new algorithm for spatially stratified differential expression, we tested >2 million hypotheses about cell types’ behavior across space. We measured how CD8^+^ T cells change upon entering the tumor, how cancer cells adapt to nutrient-poor microenvironments, how tumor glycolysis impacts nearby healthy cells, and how low-proliferation zones of the tumor are distinct. We found several instances of druggable biology: the tumor’s heterogeneity and invasion programs suggested aggressiveness independent of traditional grading; PDCD1 expression and exhaustion limited T-cell activity; cancer cells engaged angiogenic signaling but struggled to proliferate in hypoxic regions; and a tumor subcluster up-regulated lipid metabolism. These findings demonstrate the insights obtainable through spatial profiling of individual tumors.

## Introduction

Cancer is a complex system in which the microenvironment imposes evolutionary pressures, and cancer cells adapt to overcome them. Hallmarks of these adaptations include genetic plasticity, evasion of immune attack, sustained proliferation even as the microenvironment becomes depleted of nutrients, and phenotypic changes to allow growth in new milieu^1^. While these hallmarks pertain broadly across cancer, every tumor is unique, with distinct mutations and distinct tumor-microenvironment interactions driving distinct response to therapy. While DNA sequencing can measure a tumor’s mutation profile comprehensively, biomarkers of tumor and microenvironment phenotype remain coarse, relying e.g. on low-plex protein stains or bulk gene expression profiles that fail to capture the spatial context driving therapy resistance and immune evasion.

New technologies open new windows on biology. Recent high-plex spatial transcriptomics assays can now uncover not only the locations of cell types but also the spatial dynamics of cell states. Here we present the first ever breast invasive ductal carcinoma measured with whole transcriptome spatial imaging. Using this data, we sought to assess: what can spatial transcriptomics reveal about one patient’s disease, and how might these findings guide treatment?

Below we explore how the hallmarks of cancer manifest in this tumor, detailing at whole-transcriptome scale how tumor, immune and stroma cells interact across changing microenvironments. We characterize cancer cells’ phenotypic diversity across the span of the tissue, study the changing activity of T cells as they infiltrate the tumor, characterize how tumor cells adapt to lower nutrient availability, measure how cancer cells impact their neighbors through glycolysis, and use regions of reduced cancer cell proliferation to identify stressors on the tumor. Throughout, we recapitulate known biology and highlight unexpected discoveries, some with implications for treatment.

## Results

### Landscape of a tumor: morphological, microenvironment and phenotypic diversity

Taking a 1.6 cm^2^ grade II HER2^+^ / PR- / weakly ER^+^ invasive ductal carcinoma, we profiled 18935 genes across 611937 cells using the CosMx® Human Whole Transcriptome Panel. For partial validation of RNA findings, we imaged 64 proteins in a non-adjacent section from the same tumor using the CosMx Human Immuno-Oncology Protein Panel (Methods).

We began with a high-level survey of the tumor’s spatial organization. This initial review of immunofluorescent staining revealed striking heterogeneity, with regions differing in both cellular composition and structural arrangement. We noted two lobes with distinct morphology, one with densely-packed tumor glands and the other interdigitated with broad stromal regions. Between these lobes is a stroma-dominated region containing non-contiguous tumor glands (Fig 1a). In the protein slide, the dense lobe is clearly identifiable, while the 2-dimensional patterns of the interdigitated lobe and stromal regions have shifted (Supplemental Fig 1). Building on these observations, we next sought a systematic way to capture this complexity. We used the transcriptomic data to partition the tumor into nine spatial domains (Methods). These domains were smooth and fine-grained, with some domains occupying regions just 1-2 cells wide (Fig 1b, Supplemental Fig 2). They align to distinct anatomical features, from the exterior stroma (Exterior stroma 1; ES1), to the adjacent stroma surrounding the tumor (Exterior stroma 2; ES2), to the tumor interior (Tumor core 1, 2, and 3; TC1, 2, and 3), and to stroma intermingled with the tumor (Interior stroma 1, 2, and 3; IS1, 2, and 3) and its border with the surrounding tumor (Interior stroma 4; IS4).

**Figure 1.**
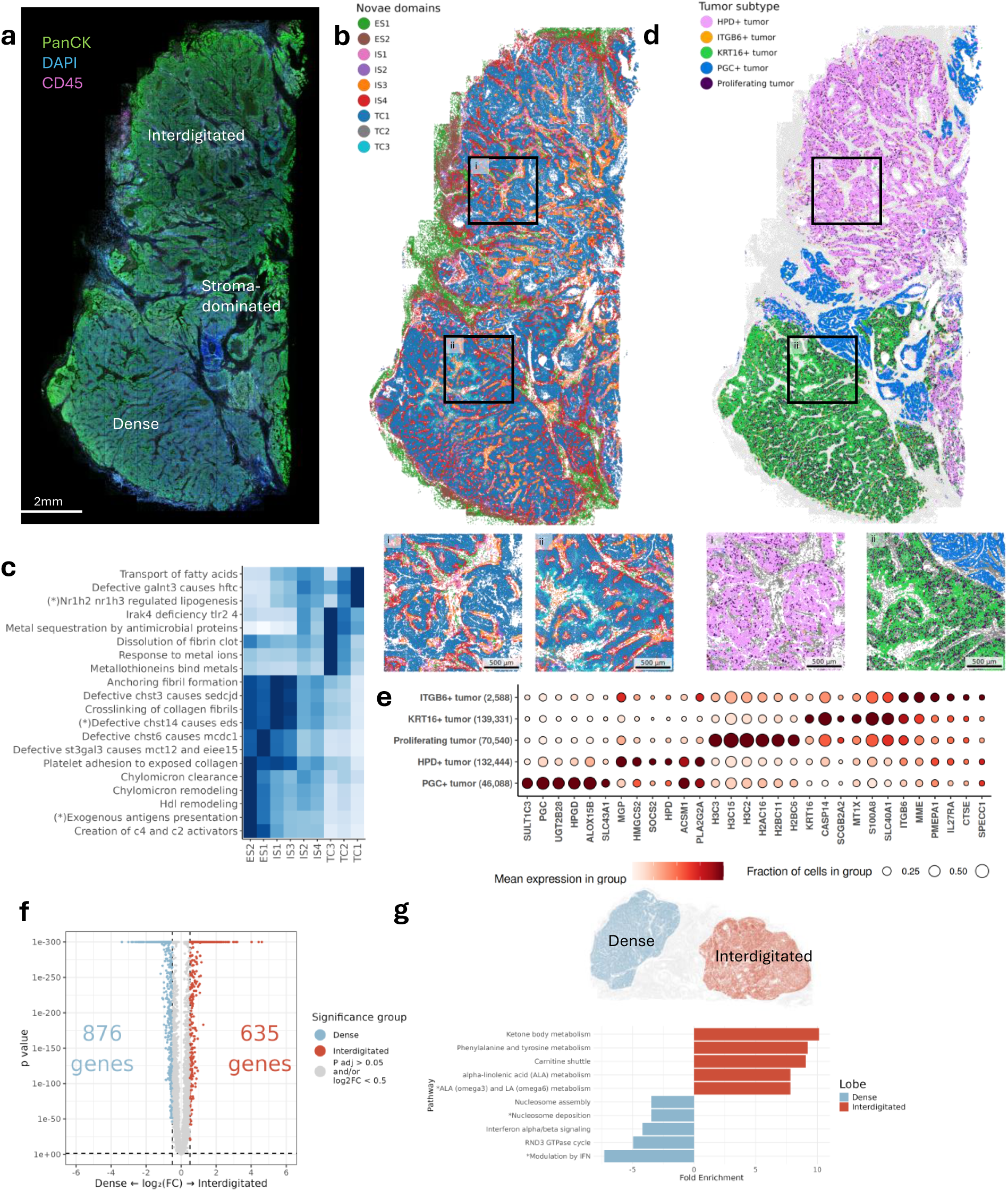
Landscape of the tumor. **a.** Immunofluorescence image of the tumor, with DNA in blue and PanCK in green. **b.** Spatial domains derived from gene expression with two insets showing zoomed in plots. **c.** Enrichment of REACTOME pathway scores (AUCell) in each spatial domain. Darker blues indicate higher scaled scores. Abbreviated pathway names are indicated by stars: “HR1H2-NR1H3-regulate-gene-expression-linked-to-lipogenesis”, “defective-CHST14-causes-EDS-musculocontractural-type”, “cross-presentation-of-particulate-exogenous-antigens-phagosomes”. **d.** Spatial distribution of cancer subclusters. Non-cancer cells are grey. Two insets show zoomed in plots. **e.** Cancer subclusters’ expression of marker genes. **f.** Differential expression results comparing the interdigitated lobe to the dense lobe. **g.** Pathway enrichment in the two lobes. Abbreviated pathway names are indicated by stars: “alpha-linolenic (omega3) and linoleic (omega6) acid metabolism", "Deposition of new CENPA-containing nucleosomes at the centromere", "Modulation of host responses by IFN-stimulated genes."

To understand the molecular differences between domains, we identified the domains’ most characteristic pathways, much as one would identify cell type marker genes (Fig 1c). Comparing the stromal interface (ES2) to the more exterior stroma (ES1), we see higher activity of antigen presentation, PD-1 signaling, and HDL remodeling, suggesting increased immune reactivity at the tumor’s border. The dominant tumor-enriched domain, TC1, has prominent metabolic signaling enrichment: transport of fatty acids, gene expression linked to lipogenesis, and activation of metabolic gene expression. The relatively rare tumor-enriched domains TC2 and TC3 are distinguished by innate inflammation (TLR/DAMP) and metal sequestration (metallothioneins and “metal sequestration by antimicrobial proteins”, a proxy for S100A8/A9-calprotectin biology). Interior stroma domains IS1 and IS2 show high levels of ECM related pathways such as crosslinking of collagen fibrils and collagen chain trimerization, suggesting a highly fibrotic region typical of vasculature and cancer-associated-fibroblast (CAF) presence. The cumulative picture that emerges is a tumor surrounded by, infiltrated by, and interacting with immune-rich stroma. Variability within tumor regions is driven in part by metabolism, innate immunity and metal sequestration. Variability within the tumor-interdigitated stroma is driven in part by CAF activity and access to vasculature.

To obtain cell types, we used a hybrid of the HieraType algorithm^2^ to finely classify immune cells and iterative Leiden clustering to identify the remaining cell types (Methods; Supplementary Fig 3). Based on spatial location, expression of epithelial markers, and high total mRNA content, we classified five of the Leiden clusters as cancerous. The cancer clusters were distinguished by strong overexpression of certain genes: KRT16, PGC, ITGB6, HPD, and proliferation markers, namely histone genes (Fig 1d,e). Some tumor cell types were strongly spatially segregated: KRT16^+^ tumor dominated the dense lobe, HPD^+^ tumor the interdigitated lobe, and PGC^+^ tumor formed glands in the stroma-rich region between the tumor’s two main lobes. The proliferating tumor cluster (18% of tumor cells) was scattered diffusely in the interdigitated lobe and more densely in the dense lobe, where they tended to fall in the interior rather than the border of tumor glands. ITGB6^+^ tumor cells were also diffuse and even more rare (0.7% of tumor cells).

The top markers that differentiated these tumor subtypes reveal that these subtypes represent distinct biological states or microenvironmental niches. The most abundant subtype, KRT16^+^, was marked by elevated levels of the stress response- and inflammation-related gene S100A8, which has been used as a biomarker for poor prognosis and relapse^3^.

The additional high expression of KRT16 points towards keratinization, often seen at the invasive and metastasizing front of tumors^4^. Densely intermingled with this subtype was the proliferating subtype, which had high expression of histones, suggesting active chromatin remodeling; this suggests the lower dense lobe was a fast-growing, aggressive tumor. In contrast, the upper, interdigitated lobe was characterized by high expression of the metabolism related genes ACSM1, HMGCS2, and HPD, reflecting a metabolically active but potentially less aggressive lobe. The PCG^+^ tumor forms lobes in the center of this section with distinct phenotypes, with comparatively higher expression of lipid and steroid signaling genes ALOX15B, UGT2B28 and SULT1C3. These genes are typically more highly expressed in differentiated epithelial cells, suggesting this was the least proliferative and aggressive region of the tumor. Finally, the rarest subtype we identified, ITGB6^+^, was marked by upregulation of invasion and EMT-related genes ITGB6, MME, and PMEPA1 in addition to immune-regulating genes IL27RA and CTSE, implicating this rare population in aggressive, leading-edge invasion.

For an orthogonal exploration of the tumor’s evolution, we compared cancer cell gene expression across the dense and interdigitated lobes using the smiDE package (Fig 1f, Supplementary Table 1). The expression of 1511 genes was significantly different with adjusted p-value <0.05 and a fold change of at least 0.5 with similar numbers of genes upregulated in each lobe. The interdigitated lobe was highly enriched (fold change > 3) for the well-known metabolical signaling genes FABP7, HMGCS2, and FASN, which likely reflect metabolic adaptations that support tumor growth and therapy resistance.

Additional highly enriched genes contribute to immune signaling, including CXCL13 and PLA2G2A. Somewhat surprisingly, this lobe also showed elevated expression of ALOX15B, a tumor suppressor associated with reduced proliferation and invasion in breast cancer^5^ suggesting this lobe may be more susceptible to chemotherapy. In comparison, the dense lobe was highly enriched for the immune and interferon related genes ISG15, IFI6, and IFTIM3. Genes known to be oncogenes and correlated with poor prognosis or aggression were elevated, including FAM83A, CRABP2, and CYP4Z1^6–8^, together pointing towards oncogenic reprogramming and therapy resistance. Consistent with these highlighted genes, gene set enrichment analysis results were dominated by metabolic pathways in the interdigitated lobe, and by interferon and nucleosome pathways in the lower lobe (Fig 1g). Together, these patterns reveal a highly heterogeneous tissue landscape, where metabolic adaptation and immune signaling coexist in separate compartments, shaping tumor evolution.

### Mechanisms of immune evasion

T cells are central to anti-tumor immunity, and their failure to clear cancer often reflects mechanisms of immune evasion that can inform therapeutic strategies. This tumor harbored a robust T-cell infiltrate, consisting of 7.3% of cells in the sample, and infiltrating throughout the tumor. As evidenced by the fact that the tumor existed to be resected, these abundant T cells had failed to eradicate it. To understand this tumor’s mechanisms of immune evasion, we characterized its interactions with its T-cell infiltrate.

The HieraType algorithm identified 23 immune populations, including 12 T-cell subsets, displaying distinct expression of canonical marker genes (Fig 2a). T cells were found throughout the tumor, clustering densely in both the exterior stroma and interior stroma, and appearing more sporadically in the tumor bed (Fig 2b,c). Protein profiling confirmed that T cells were densely packed at the outskirts of the dense lobe and infiltrating at lower frequencies within it, that CD8^+^ T-cells infiltrates into tumor glands at much higher rates than CD4^+^ T cells (Supplemental Fig 4), and that naïve (CD45RA^+^) T cells seldom entered the interior stroma or tumor interior (Supplemental Fig 5). T cells infiltrated into the tumor bed more often in the dense lobe than in the interdigitated lobe. CD8^+^ T-effector memory cells were abundant in the exterior stroma and found infrequently in the tumor bed.

**Figure 2.**
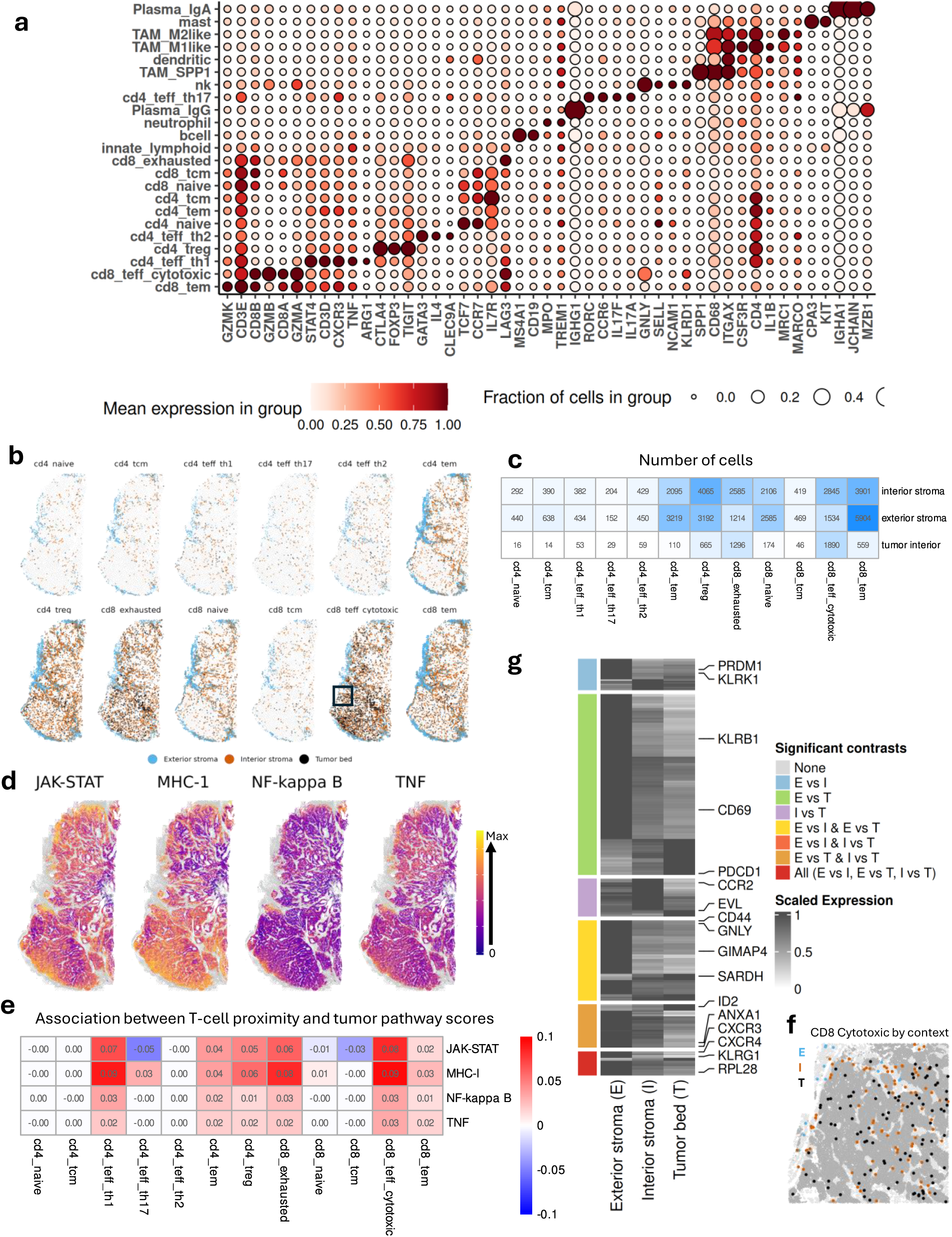
Cancer cell interactions with the T-cell infiltrate. **a.** Immune subcluster expression of marker genes. **b.** Locations and spatial contexts of T-cell subpopulations. **c.** Abundance of T-cell subpopulations across spatial contexts. **d.** Spatially smoothed pathway scores. Only cancer cells are colored. **e.** Association between cancer cells’ pathway scores and abundance of T-cell subpopulations in their 100 nearest neighbors. Color denotes the increase in z-transformed pathway score associated with each additional T cell in their nearest neighbors. Color only given to associations with p < 0.01. **f.** Relative expression of differentially expressed genes in CD8 cytotoxic effector T cells across spatial contexts. All genes reaching significance in any pairwise comparison (FDR < 0.05) are shown; genes of interest are labeled. **g.** Zoomed inset from the region boxed in (b) that shows the three spatial contexts for CD8 cytotoxic T cells. Tumor cells in dark grey and non-tumor cells in light grey.

Regulatory T cells (Treg) had a similar distribution. In contrast, exhausted and cytotoxic CD8^+^ T cells were most abundant in the interior stroma and were also present at high rates in the tumor bed. Thus at the moment this tumor was removed, 48% of the CD8^+^ T cells in the tumor bed were cytotoxic and not yet exhausted. Although these cytotoxic CD8^+^ T cells represented just 0.6% of the tumor bed’s census, they nonetheless had the theoretical potential to eliminate dozens of tumor cells each. With more T cells presumably trafficking to the tumor continually, the tumor would need some means of disabling them.

A successful T cell encounter with a tumor cell kicks off multiple causal chains. In one, T-cell interferon signaling stimulates the JAK-STAT pathway, which in turn induces MHC-I antigen presentation^9^. In another, TNF signaling induces apoptosis. The cancer cell can resist this attack through pro-survival NF-kB signaling and presentation of immune checkpoints^10^. Plotting REACTOME pathway scores for these processes across the tissue’s span specifically in tumor cells, we see JAK-STAT and MHC-I antigen presentation throughout much of the tumor, concentrated in regions dense in cytotoxic CD8^+^ T cells (Fig 2d). The two lobes differ in these T-cell related pathways: the dense lobe has ubiquitously high MHC-I antigen presentation, while the interdigitated lobe only up-regulates this pathway when in direct contact with the stroma. Meanwhile JAK-STAT signaling is more uniformly high in the interdigitated lobe. TNF and NK-kappa B signaling are high mainly at the tumor’s periphery. To further investigate the impact of T-cell subpopulations on tumor cells, we ran multivariate models predicting tumor cells’ pathway scores from the abundance of T-cell populations in their 100 nearest neighbors (Fig 2e). These 4 pathways were most strongly associated with presence of cytotoxic and exhausted CD8^+^ T cells and Th1 effector CD4^+^ cells, and to a lesser extent with Tregs and CD4^+^ and CD8^+^ effector memory cells. MHC-I and JAK-STAT responded 2-3x more strongly to T-cell presence than NK-kappa-B and TNF did.

Thus the T-cell infiltrate is seen to perform its expected functions, and the tumor appears to employ NF-kappa B to defend against T-cell attack. However, because NK-kappa B pathway expression is so concentrated at the edge of the tumor, it only partially explains this tumor’s robustness to infiltrating T cells.

To further investigate the sources of the tumor’s survival, we examined the expression patterns of cytotoxic CD8^+^ T cells as they infiltrated into the tumor. Specifically, we grouped these T cells into 3 spatial contexts: exterior stroma (novae niches ES1 and ES2), interior stroma (niches IS1, IS2, and IS3) and tumor interior (niches TC1, TC2, TC3, and TC4) (Fig 2f). We then used the smiDE package to perform differential expression analysis of cytotoxic CD8^+^ T cells across these three locations (Fig 2g, Supplementary Table 2). smiDE’s overlap ratio filter, which flags genes at risk of bias from cell segmentation errors, identified 8954 genes as safe to analyze in these T cells. Of these genes, 435 produced an adjusted p-value (Benjamini-Hochberg < 0.05) in at least one pairwise comparison between spatial contexts.

Contrasting cytotoxic CD8 cells in tumor interior with those in the exterior stroma, we find tumor-interior CD8 cells to up-regulate checkpoint and exhaustion genes PDCD1, ID2 and SARDH^11^. They down-regulate the effector genes KLRK1, KLRG1, GNLY and PRDM1, the homing and adhesion gene CD44, the chemotaxis genes CXCR4 and SELPLG, and TXNIP, which holds T-cell metabolism in check until released by TNF signaling^12^. More unexpectedly, they also down-regulate GIMAP4, a relatively under-investigated gene which has been linked to control of cell death and cytokine secretion in T cells^13^. This loss of GIMAP4 expression while in the antigen-rich and exhaustion-promoting tumor interior is an unexplored potential mechanism of tumor-driven T-cell inhibition. Finally, cytotoxic CD8^+^ cells in the tumor interior down-regulated the ribosomal genes RPL28, RPL13, RPL17, and RPS20, suggesting a loss of translational activity consistent with an increasingly exhausted state. The cumulative picture is of cytotoxic CD8^+^ cells infiltrating the tumor, then losing effector, homing and translational activity and initiating exhaustion programs. Several unexpected genes emerged from this analysis, including AKAP5, IFITM10, DGHK and SARDH, which were up-regulated in the tumor bed; and TRDC, NBL1, and EMP3, which were down-regulated.

The interior stroma presented as a transitional niche, with increased expression of CCR2 and EVL indicating active migration and adaptation, and with upregulation of histone genes (H2BC4, H1-4) pointing to epigenetic remodeling.

### Cancer cells’ adaptations to unavailable vasculature include angiogenesis, glycolysis, and up-regulated ELF3

Cancer cells face the challenge of continuing proliferation even as they collectively consume available nutrients at high rates. To study how cancer cells adapt to nutrient-poor microenvironments, we measured how their gene expression changed as they grew further from blood vessels. Blood vessels were defined as any collection of at least 20 vascular cells within a 0.1 mm radius, as identified with dbscan (Methods) (Fig 3a). Protein profiling of the dense lobe confirmed the general pattern of vasculature identified by mRNA

**Figure 3.**
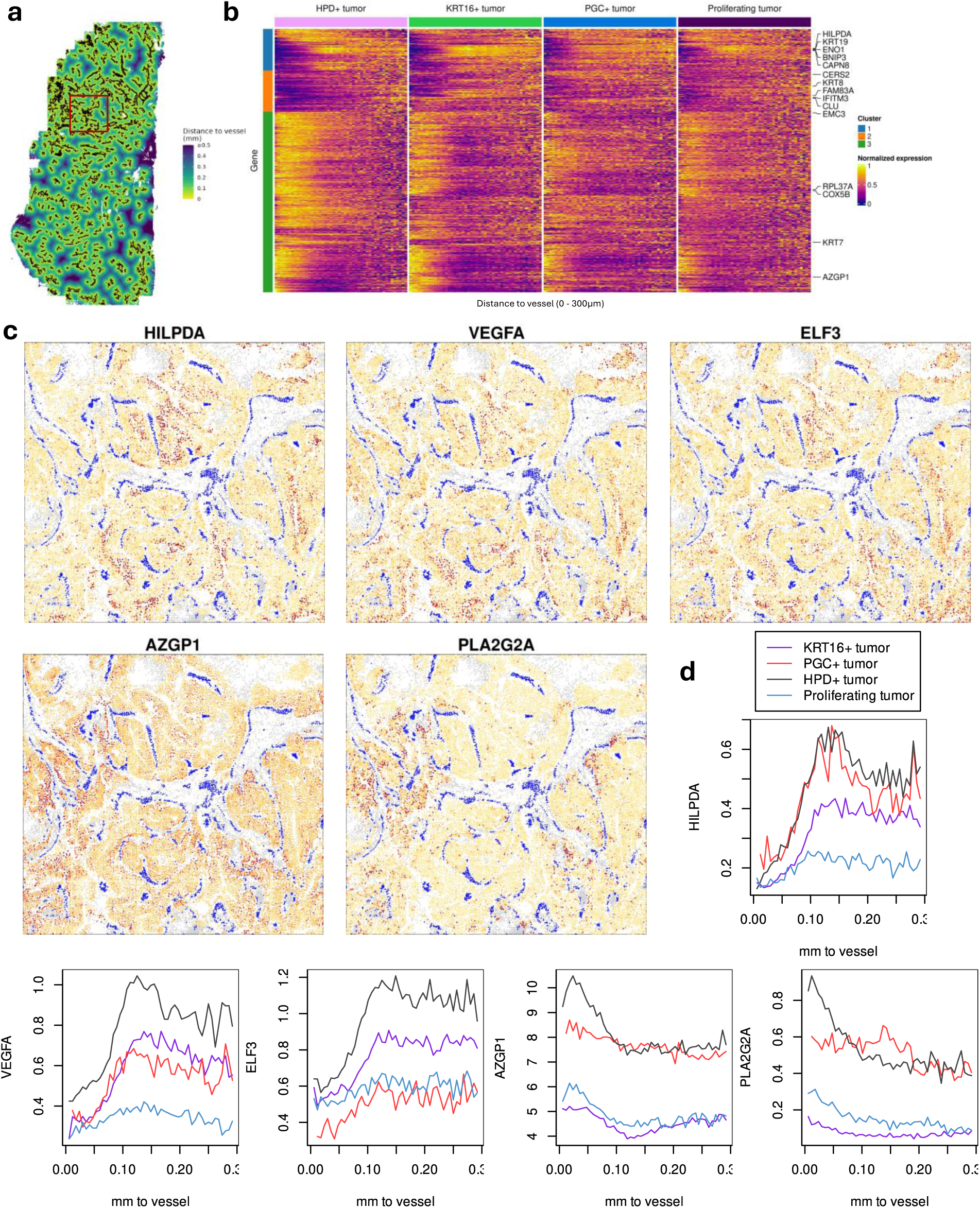
Cancer cell expression changes with proximity to vasculature. **a.** Spatial plot of vasculature (black) and all other cells colored by distance to vasculature. **b.** Differential expression in cancer cells vs. distance to vasculature. For each of 4 cancer subtypes, mean expression across 49 distance bins is shown. The 1000 genes with smallest p-values are show; genes of interest are labeled. **c.** Spatial plots of selected genes. Vasculature is shown in blue, cancer cells are colored by expression level (darker = higher), and other cell types are grey. Coordinates match the orange overlay box in panel (a). **d.** Cancer cells’ mean expression of selected genes across 49 bins of distance to vasculature.

(Supplemental Fig 6). We then tested each cancer subtype separately for differential expression with respect to distance to blood vessels (Methods). Of the 13290 genes analyzed in at least one tumor subtype (genes at risk of cell overlap errors were omitted on a per cell type basis and so may not have been evaluated in each cell type), 3348 attained a BH adjusted p-value < 0.001 in at least one subtype (Supplementary Table 3). Selecting the top 1000 genes by average adjusted p-value across tumor subclusters, we found diverse expression patterns across cancer subtypes and across distances to vessels (Fig 3b). Due to low cell numbers we excluded the rarest tumor subtype, ITGB6^+^ tumor, from this visualization. Clustering these genes by their trends with distance to blood vessels, we identified 3 distinct groups. The largest cluster is up-regulated near blood vessels, then loses expression sharply around ∼0.1 mm away from vessels. Gene set enrichment analysis of this cluster found strong (adjusted p < 0.05) enrichment of the following pathways, among others: “regulation of expression of SLITs and ROBOs”, “eukaryotic translation elongation” and “Cap-dependent translation-initiation”, together suggesting a highly active cellular state with enhanced and regulated protein production.

Two other clusters of genes were enriched further from vasculature, though at varying distances. These clusters had high enrichment of REACTOME pathways “gluconeogenesis”, “glycolysis”, “metabolism of carbohydrates and carbohydrate derivatives”, and “neutrophil granulation”. Together, these paint a picture of enhanced glucose metabolism and certain immune interactions further from vessels.

Some genes with notably strong results deserve mention (Fig 3c,d). HILPDA was the gene most strongly up-regulated in cancer cells distant from blood vessels; this gene is a known component of cells’ hypoxia response^14^. In the interdigitated lobe of the tumor, HILPDA was often most strongly up-regulated on the interior wall of tumor glands, marking these structures as oxygen-deprived compared to the less structured cancer cells spreading from them. VEGFA was also strongly up-regulated away from vasculature. Its expression climbed sharply starting ∼0.05 mm from vessels, peaked around 0.12-0.15 mm from vessels, and declined slightly beyond there. Proliferating tumor cells expressed VEGFA at much lower levels that other tumor subtypes. More unexpectedly, ELF3 was also strongly up-regulated away from vessels. ELF3 is a transcription factor controlling differentiation that has been associated with poor outcomes in HER2^+^ breast cancers^15,16^. AZGP1 was the gene most up-regulated near blood vessels, with expression dropping quickly until bottoming out ∼0.1 mm from vessels. AZGP1 promotes lipolysis, suggesting that the cancer cells near vessels were taking advantage of a metabolic pathway unavailable further away.

Also strongly up-regulated near blood vessels was PLA2G2A, which catalyzes the hydrolysis of phospholipids in cell membranes^17^. This gene was only activated in the interdigitated lobe of the tumor. Most cells with CD31^+^ protein stains were also CD34^+^ (Supplemental Fig 6); the rarity of CD31^+^/CD34^-^ cells suggests actively developing vasculature, consistent with widespread angiogenesis. The combination of active vascular remodeling and hypoxia-driven gene expression points to an adaptive system in which cancer cells exploit both proximity to vessels and metabolic rewiring to keep proliferating as conditions change.

### Impacts of glycolysis on the tumor microenvironment

Tumors employ glycolysis to rapidly generate ATP and to overcome hypoxic microenvironments. Glycolytic cells simultaneously take glucose from the microenvironment and secrete lactate, which acidifies the microenvironment. Both these effects act as stressors on nearby cells^18,19^. This spatial, single-cell and whole transcriptome dataset presents an opportunity to examine *in vivo* the transcriptome-wide impacts of glycolysis on diverse cell populations.

To measure cells’ exposure to the by-products of glycolysis, we first used AUCell to score every cell for activity of the REACTOME glycolysis pathway. Then, we averaged these glycolysis scores over each cell’s 50 nearest neighbors, producing a metric of each cell’s “neighborhood glycolysis” (Fig 4a). Cancer cells were the dominant contributor to cells’ neighborhood glycolysis scores, contributing 72.7% of the tissue’s total glycolysis scores.

**Figure 4.**
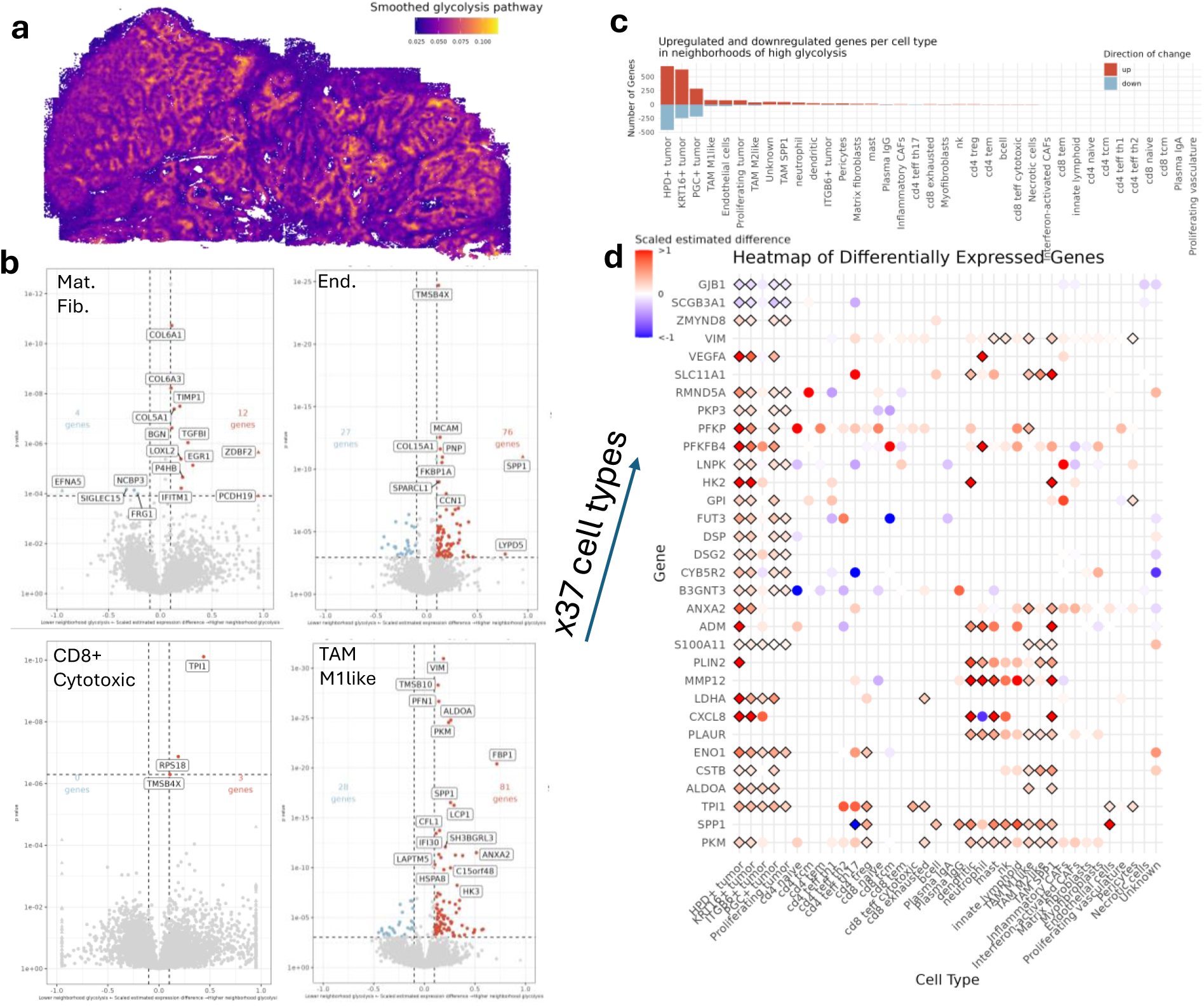
Neighborhood glycolysis levels drive concerted transcriptional changes. **a.** Spatially smoothed glycolysis pathway score. **b.** Differential expression between high and low glycolysis neighborhoods in selected cell types. X-axis corresponds to scaled expression change, with red indicating higher expression in high glycolysis neighborhoods and blue indicating reduced expression. Scaled expression difference was capped at +/-0.95 and capped data points are shown as triangles. **c.** Number of differentially expressed genes per cell type. **d.** Scaled difference in gene expression between high and low glycolysis neighborhoods for all cell types and genes reaching significance threshold in at least four cell types. Genes with significant differences (FDR < 0.05) shown as diamonds.

We then performed a series of differential expression analyses, modeling how each cell type’s expression profile changed as its neighborhood glycolysis increased, beyond effects from each cell’s spatial domain and tumor neighbors (Methods) (Supplementary Table 4).

Of particular interest are T cells, which are suppressed in glycolytic environments^18^; macrophages, which polarize to a more inflammatory state; fibroblasts, in which glycolysis activates ECM remodeling and excludes immune cells^20^; and endothelial cells, which are spurred to angiogenesis^21^. Our results support and extend these prior findings (Fig 4b).

Within endothelial cells, regions of high glycolysis showed a shift toward an angiogenic and stress-adapted phenotype. These glycolytic regions down-regulated ELN, CLEC3B, and PTPRB, which are associated with vascular elasticity, extracellular matrix remodeling, and maintaining vascular junctions, respectively^22–24^. Also down-regulated were IFI44L and IFIT1, suggesting dampened interferon-driven inflammation in high glycolysis regions.

Conversely, LDHB, PGAM1, and HSPA5 were all elevated, consistent with enhanced glycolytic flux and ER stress adaptation, alongside increased SPP1, CD276, and JAK1, pointing to immune modulation and pro-angiogenic inflammation. Up-regulation of ROBO4, THBS1, and TIMP1 further reflects dynamic remodeling of the vasculature under metabolic stress.

Contrasting the expression patterns of M1-like macrophages across regions of high and low neighborhood glycolysis revealed a shift towards a metabolically active, pro-inflammatory M1 phenotype in glycolytic zones. In regions of high glycolysis, M1-like tumor-associated macrophages (TAMs) upregulated immune activation markers such as MYD88, CD44, and SPP1 as well as glycolytic enzymes such as PKM, PFKP, and ALDOA, consistent with elevated inflammatory signaling and energy production. In contrast, in regions of low glycolysis, the M1-like TAMs exhibited evidence of a shift towards a more M2-like state, with upregulation of the M2-associated markers CD209, FOLR2, and C1QA, suggesting a more immunosuppressive state.

Interestingly, cytotoxic CD8 T cells showed minimal expression changes between high and low glycolysis regions, with only 3 genes up-regulated with FDR < 0.05 in glycolytic zones: TPI1, a glycolytic enzyme; RPS18, which suggests elevated translational activity; and TMSB4X, which involved in trafficking. Together, these genes are suggestive of active T cells.

In regions of high glycolysis, matrix fibroblasts exhibited a transcriptional profile consistent with ECM remodeling and tumor-supportive activity. Upregulated genes included collagens, LOXL2, TGFBI, and TIMP1, all of which contribute to ECM stiffness and remodeling^25–27^. Conversely, a handful of genes were downregulated in high glycolysis neighborhoods including the immune checkpoint gene SIGLEC15 and the cell-cell communication gene EFNA5, suggesting a shift in immune signaling.

We also sought genes whose response to glycolysis was consistent across cell types. Collating the differential expression results from each cell type (Fig 4c), we identified 32 genes that were significant at FDR < 0.05 and had a scaled effect greater than 10% in at least four cell types (Fig 4d). Of these genes, 30 were up-regulated in glycolytic regions, while only 2 were consistently down-regulated. Frequently up-regulated genes in glycolytic neighborhoods included PKM, SPP1, TPI1, ALDOA, CSTB, ENO1, and PLAUR. Summarizing trends across multiple cell types, we find that regions of high neighborhood glycolysis were characterized by the coordinated upregulation of genes involved in metabolic adaptation and immune modulation. In particular, PKM, a glycolytic enzyme important in enhancing glycolytic flux and a hallmark of the Warburg effect^28^, was significantly upregulated in almost a third of the cell subtypes. The next most frequently upregulated gene, SPP1, is a multifunctional mediator that promotes immune evasion and stromal activation, consistent with a pro-tumoral microenvironment. PLAUR and MMP12 were significantly up-regulated across diverse myeloid cell types. PLAUR assists cell migration and adhesion^29^, while MMP12 can assist invasion by degrading the ECM^30^. Together, these changes suggest that these glycolytic niches within the breast cancer section were not only metabolically specialized but also structurally and immunologically adapted to support tumor progression.

### Regions of reduced proliferation suggest stressors on the tumor

Sustained proliferation is a defining hallmark of cancer, yet tumors are not uniformly proliferative. We therefore hypothesized that low-proliferation neighborhoods, where the tumor failed to thrive, might reveal stressors on cancer cells that could motivate future therapeutics. To this end, we compared cancer cell expression in low-proliferation vs. high-proliferation neighborhoods. The cell cycle phase for all cancer cells was classified using Seurat. 35.6% were in G1 phase, corresponding to resting in this framework, while 40.0% were in S phase and 24.4% were in G2/M phase. This high rate of active proliferation extended across most of the tumor; however, 10.7% of cancer cells fell in low-proliferation neighborhoods, defined as cellular neighborhoods where ≥50% of cancer cells were in G1 phase (Fig 5a, Supplemental Fig 7). Protein Ki67 stain confirmed the large low-proliferation region along one edge of the dense lobe (Supplemental Fig 8).

**Figure 5.**
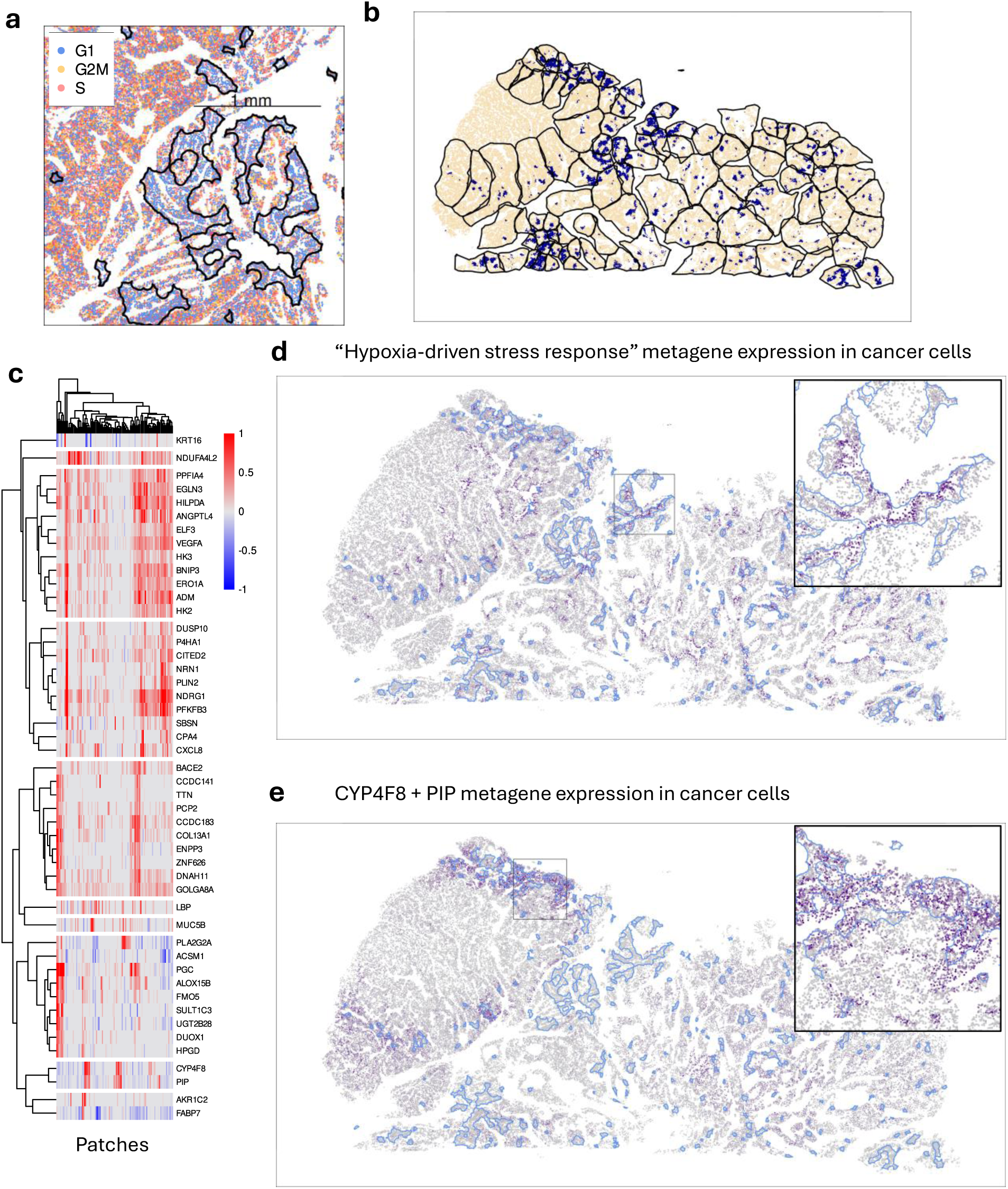
characteristics of cancer cells in low-proliferation neighborhoods. **a.** Definition of “low proliferation neighborhoods”: ≥50% of neighboring cancer cells in G1 phase. Polygons highlight cells meeting this criterion. Only cancer cells are plotted. **b.** Patches for stratified differential expression analysis. Cells within low-proliferation neighborhoods are in dark blue; patch borders are in black. Only G1-phase cancer cells are plotted. **c.** Differential expression results for selected genes. Patches are in columns; genes with strong results in at least 3 patches are in rows. Color shows effect size (proportion change in mean expression when in low-proliferation neighborhoods); results with p > 0.05 are left grey. **d, e.** Expression of the “hypoxia-driven stress response” and “CYP4F8+PIP” metagenes. Low-proliferation neighborhoods are outlined in blue. Only G1-phase cancer cells are shown.

Low-proliferation neighborhoods fell in diverse microenvironments, leading us to assume that these cancer cells might be encountering different stressors in different patches. This motivated a novel approach to differential expression analysis: rather than modeling all cells at once, we split the tumor into dozens of small patches, then ran a separate differential expression analysis within each patch (Fig 5b). Patches were selected with a new algorithm designed to maximize statistical power within each patch while constraining patch size and shape (Methods). To avoid confounding from cell cycle phase, we confined this analysis to cells classified as G1, or “resting” phase.

We then ran differential expression models separately within each patch, comparing the G1 tumor cells within high and low proliferation neighborhoods. We confined our analysis to 18141 genes determined by smiDE to be at low risk of bias from cell segmentation errors. This produced a grid of DE results for 18141 genes across 100 patches, allowing discovery of both global and local correlates of tumor proliferation (Supplementary Table 5).

In hopes of isolating factors with more causally direct links to proliferation, we selected genes with highly significant and strong effect sizes (p < 0.01, effect size > 50% of the gene’s mean expression) in at least 3 patches. This filter returned 48 genes. Confirming the hypothesis that suppressed proliferation has diverse origins, none of these genes had ubiquitous findings across the span of the tumor. Clustering genes based on their differential expression results across patches (Fig 5c), we find three consistent themes. First, a cluster of genes involved in hypoxia-driven stress response (ADM, ANGPTL4, BNIP3, EGLN3, ELF3, ERO1A, HILPDA, HK2, HK3, PPFIA4, VEGFA) was up-regulated in a large share of low-proliferation neighborhoods across the tumor (Fig 5d; p<0.01 in 35-68% of patches). This cluster’s expression often extended beyond the core low-proliferation neighborhoods, suggesting a model in which cancer cells can sustain proliferation in the presence of moderate hypoxia but fail to compensate in more intensely hypoxic zones.

Many of these genes were also identified as differentially expressed as a function of distance to the nearest vasculature, consistent with this representing a hypoxic signature. A second cluster, PIP and CYP4F8, marked low-proliferation regions found along the edges of only the dense lobe. These genes have more cryptic relationships to low proliferation: CYP4F8, which catalyzes production of prostaglandin^31^, has been identified as a drug target for prostate cancer^32^; while PIP, typically associated with luminal and apocrine breast cancer subtypes, has been linked both to high^33^ and low proliferation^34^. A third cluster of genes (ACSM1, ALOX15B, DUOX1, FMO5, HPGD, PGC, PLA2G2A, SULT1C3, UGT2B28) corresponded to regions of PGC^+^ tumor.

## Discussion

The sample analyzed above is the first breast tumor and one of the first tumors of any kind to be profiled via spatial imaging of the whole transcriptome. We found that this single experiment provided a high-resolution view of this sample without support from duplicate assays. We resolved 37 cell types, including 12 T-cell subpopulations and 11 other immune cell types, obtaining distinct profiles of canonical marker genes. We also obtained clear differential expression results even from low-expressing genes, e.g. PDCD1 in cytotoxic CD8 T cells. The scale of the results is also notable: we tested 8066 differential expression hypotheses in Fig 1, 8954 in Fig 2, 24013 in Fig 3, 310823 in Fig 4, and 1814100 in Fig 5.

Most questions we pursued culminated in a differential expression analysis, asking variants of the question, “how does a given cell type change in response to some attribute of its neighborhood?” This approach led to conclusions about how two morphologically distinct lobes of the tumor differed, how cytotoxic CD8 T cells changed as they invade tumor glands, how tumor cells reacted as they fall farther from blood vessels, how diverse cell types were impacted by local glycolytic activity, and how tumor cells in regions of low proliferation were distinct.

This kind of differential expression analysis is only possible with data that is both single cell and spatially resolved. Spot-based assays would be unable to support differential expression of low-density cells like CD8 T cells, as their expression profiles would be too intermixed with those of surrounding cells. Meanwhile, dissociated single cell assays would be blind to most of the predictor variables we queried. The richness of the answers we obtained depended linearly on the plex of the assay: measuring half as many genes would have yielded half as many findings.

Our analyses were limited mostly by the questions we chose to pursue, and constitute a tiny fraction of what could be learned from this dataset. For example, given our 37 cell types and 18935 genes, analyses of the form “how does cell type X change with distance to cell type Y” would support 1369 differential expression analyses and >25 million hypothesis tests. A full survey of this and other differential expression configurations would beggar human comprehension, with two implications. First, given a distinctive biological question, future analysts will be able to find noteworthy results from re-analysis of existing datasets like this one. Second, as more datasets of this richness are published, the biology contained within them may be more comprehensively “understood” by computer resources like foundation models, large language models, or simply databases of correlations.

To encourage the best use of this data type in future studies, we sought throughout to employ optimal and novel analysis strategies. Notable tools include the hieraType algorithm^2^ for nuanced immune cell typing, the smiDE package’s toolkit for protecting differential expression from bias arising from cell segmentation errors, and the patchDE algorithm for stratifying differential expression problems across spatial patches. This last algorithm, created for this study, proved productive, and we propose it as a promising new approach to spatial differential expression analysis.

Many of our results involve biology that could be relevant to treatment. The heterogeneity of the tumor across space suggests the presence of at least three major clones (KRT16^+^, HPD^+^, PGC^+^) forming distinct lobes. Genetically diverse tumors have greater potential to evolve treatment resistance^35^ and therefore may merit more aggressive treatment. In addition, the rare ITGB6^+^ tumor subcluster upregulated genes associated with invasion (Fig 1e), suggesting a path by which cancer cells would break free of the tumor bed. This provides an indicator of tumor aggressiveness that is distinct from grade, stage and proliferation.

Meanwhile, response to checkpoint inhibition has been linked to the abundance, location and expression patterns of T cells. In this tumor, cytotoxic CD8 T cells were abundant (1.0% of cells), and they successfully infiltrated from the exterior stroma to the interior stroma and to the interior of tumor glands (Fig 2c). This is consistent with an immunogenic tumor that is not structurally excluding T cells. The tumor’s expression of MHC-I and JAK-STAT pathways in proximity to T cells (Fig 2d,e) suggests these T cells were not mere bystanders but were actively engaging the tumor cells. However, upon reaching the tumor interior, cytotoxic CD8 cells up-regulated exhaustion markers, including the checkpoint gene PDCD1 (Fig 2g), a well-established dynamic that checkpoint inhibitors were designed to overcome.

Anti-angiogenic therapies have been approved for several cancer types, though not breast cancer. This tumor showed strong angiogenesis signaling, with dense vascularization (Fig 3a), a strong cancer cell response to blood vessel proximity (Fig 3b) and up-regulated VEGFA in cancer cells further from vessels (Fig 3c). Furthermore, we found a hypoxia-associated metagene enriched in regions of low proliferation (Fig 5d), implicating hypoxia as a check on this tumor’s growth.

Metabolic pathways emerged as a common motif in our results: the interdigitated lobe was characterized by up-regulated metabolic pathways (Fig 1g), and glucose metabolism was enriched further from blood vessels (Fig 3b) and in low proliferation zones (Fig 5c).

Glycolysis inhibitors are in trials, but none are yet approved. The PGC^+^ tumor cluster showed increased expression of genes involved in lipid metabolism (Fig 1e). A drug for inhibiting fatty acid synthesis, denifanstat, is currently in phase II trials in HER2^+^ breast cancer^36^.

This study illustrates how whole-transcriptome spatial imaging can expose the complexity of tumor ecosystems within intact tissue. Even in a single breast cancer specimen, we observe heterogeneity in cancer cell states, immune engagement, and metabolic adaptation—features that bulk profiling would miss. While this is an n-of-1 analysis, it demonstrates what becomes possible when spatial context is integrated with transcriptomic depth and sets the stage for broader studies aimed at uncovering principles of tumor biology that may inform treatment.

## Methods

### Data Availability

Whole transcriptome CosMx SMI data were generated from a FFPE-preserved, 5µm thick, stage II HER2^+^ breast invasive ductal carcinoma tissue sample as described in Khafizov et al 2024^37^. From a non-adjacent serial section, 64-plex protein data were generated as described below. Both of these datasets will be made publicly available on the Bruker Spatial Biology website prior to publication.

### Code Availability

All code used to run the analyses and reproduce the figures in this manuscript will be made publicly available on GitHub.

### CosMx RNA acquisition

One FFPE-preserved, 1.6cm^2^ breast tumor section was prepared for CosMx Spatial Molecular Imaging (SMI) with the CosMx Human Whole Transcriptome Panel as described in *Khafizov et al* 2024^37^. Briefly, tissue sections were mounted on Superfrost Plus slides, baked at 60 °C, and processed through deparaffinization, proteinase K digestion, and heat-induced epitope retrieval to expose target RNAs. Fiducial markers were applied for image registration, and samples were fixed and blocked prior to probe hybridization.

Whole-transcriptome (WTx) in situ hybridization probes targeting 18,935 human protein-coding genes were denatured and hybridized overnight at 37 °C. Post-hybridization washes removed unbound probes, and slides were assembled into flow cells for imaging on the CosMx SMI instrument. Prior to RNA readout, samples were stained with a panel of four antibodies (PanCK, CD298/B2M, CD68, and CD45) and DAPI for cell boundary identification. RNA detection was performed using iterative cycles of reporter hybridization and imaging, followed by fluorophore cleavage, as previously described^37^.

Image processing, transcript decoding, and cell segmentation were performed using the AtoMx® Spatial Informatics Platform, following published algorithms^37^.

### CosMx protein acquisition

A non-adjacent, FFPE-preserved section from the same breast tumor as above was processed for high-plex spatial protein profiling using the CosMx SMI with the CosMx Human Immuno-Oncology Protein Panel as described in the CosMx SMI manual^38^. After deparaffinization and antigen retrieval, the slide was incubated overnight at 4°C with a cocktail of 64 oligonucleotide-conjugated primary antibodies as well as four antibodies used for morphology plus DAPI, diluted in blocking buffer. Following antibody incubation, fiducial markers were applied for image registration, and samples were fixed in paraformaldehyde. The stained section was assembled into a flow cell and loaded onto the CosMx SMI instrument for cyclic imaging. Protein targets were detected through iterative rounds of fluorescent reporter hybridization and imaging. Raw images were processed to generate spatial maps of protein expression, and cell segmentation was performed to assign protein signals to individual cells as described previously^39^.

### Data Preprocessing

As quality control, we removed all cells with fewer than 200 or more than 15,000 transcripts, as well as all cells with an area more than five times the geometric mean of all cells’ area. 33,303 cells were flagged for removal, leaving 611,937 cells for analysis. Count data were total count normalized, 3000 highly variable genes (HVG) were identified, the normalized HVG data were scaled, 50 principal components were identified, and cells were projected into 2-D UMAP space.

### Protein image processing

Protein images were generated using Napari and the Napari-CosMx plugin^40,41^. Each protein’s contrast limits were adjusted for image clarity; no other transformations were performed. Raw protein images are available with this study’s data package.

### Spatial Domain Assignment

Functional spatial domains were identified using the deep learning model novae^42^. Spatial neighbors were identified within a radius of 80 microns and the pretrained “novae-human-0” model was fine-tuned using the CosMx SMI data with the maximum number of epochs set to 50, followed by computing representations. Multiple resolutions of domain assignment were assessed, from 1 to 20 domains, and the 9-domain assignment was chosen after visual review of domain spatial distributions and hierarchical relationships. Domains were named based on spatial localization and cell type composition.

### Cell Typing

Cell types were assigned in a two-step process composed of iterative unsupervised clustering for tumor and stromal populations and supervised hierarchical marker-based immune annotations. First, cells were divided into major clusters with Seurat’s “FindClusters” run with a resolution of 0.7 on the UMAP neighbor graph. Clusters were named on the basis of spatial localization and top gene markers. For tumor, fibroblast, and vascular clusters, cells were subjected again to preprocessing such that scaling and HVG identification could happen on a per major cell type basis, and clustering was performed again on these subsets to identify cell subtypes. For all major clusters identified as immune cells, cell types and subtypes were assigned using the hierarchical marker-based algorithm HieraType^2^. Overall, 37 minor cell types were assigned and grouped into 14 major cell types.

### Pathway Scoring

Every cell was scored for 1692 REACTOME pathways^43^ using the gene rank based algorithm AUCell^44^ with the argument ‘aucMaxRank’ set to the median number of unique genes detected per cell in the study, 1051. Top pathways per spatial domain were selected using the ‘clusterwise_foldchange_metrics’ function from HieraType^2^ followed by selection of pathways with the largest fold change for each domain relative to all other domains. When smoothed pathway scores were needed, scores were averaged across 50 nearest spatial neighbors.

### Association between pathway scores and local T cell abundance

Pathway scores were standardized to have SD = 1 across tumor cells. The number of each T-cell subpopulation within the 100 nearest neighbors of each tumor cell was recorded. For each pathway, a multivariate linear model was fit predicting pathway score from the local T-cell subpopulation counts and from the tumor cell’s subcluster (HPD^+^, PGC^+^, KRT16^+^, ITGB6^+^, proliferating).

### Vasculature Identification

Vessels were identified by selected cells with the major cell type ‘vasculature’ and applying dbscan clustering (eps = 0.1, minPts = 20) to their spatial coordinates; clusters with non-zero labels were retained as vessels for downstream analysis.

### Differential Expression

Differential expression was used to compare cells across multiple contrasts throughout this manuscript; all of these relied on the smiDE package^45^ which was designed for differential expression analysis of spatially resolved transcriptomic data. First a “preDE” step was used to enumerate spatial neighbors that may contribute to bias in gene expression at cell borders. Next, genes were restricted, on a per cell type basis, to only genes with higher expression in the cell type of interest than surrounding cells, using genes with a ratio < 1 from the “overlap_ratio_metric” function. Differential expression was calculated using the “smi_de” function. See Supplementary Table 6 for specific formula and distribution used in each figure. Pairwise or estimated mean expression per factor level were extracted from the results for plotting. Following differential expression, fgsea^46^ was used to identify enriched pathways within upregulated or downregulated gene sets.

### PatchDE Algorithm

Here we introduce the PatchDE algorithm for dividing a tissue into patches of similar statistical power to run differential expression against a variable X. PatchDE negotiates several competing desires. Often, we want patches that are reasonably circular. To limit spatial confounding within patches, they shouldn’t grow excessively large. And most importantly, patches should all have adequate statistical power, determined by their sum of squared errors in X, SSE(X).

PatchDE works as follows. Call X the predictor variable. Initial patches are defined around hotspots of variance in X. For each cell, we record the variance of X in its 100 closest spatial neighbors, use the Getis–Ord G* statistic to identify cells in variance hotspots, then use k-means clustering of hotspot cell locations to define initial patches.

Next, an iterative algorithm is used to grow and shift these patches in pursuit of more uniform SSE(X) across patches. Two quantities drive the iterations. First, each patch is given a “hunger” score, defined as the value of 1/SSE(X) calculated over its cells. Second, each cell is scored for proximity to each patch. A hybrid distance metric is defined as a weighted geometric mean of a cell’s distance to the patch center and its distance to the nearest cell in the patch; a cell’s proximity to the patch is defined as the inverse of this distance. By tuning the weighting of distance to patch center vs. distance to the nearest cell within the patch, one can force more circular patches vs. a more aggressive pursuit of uniform SSE(X). To stabilize the rate of patch changes, cells beyond user-defined distances to either the patch center or the nearest cell within the patch are given proximity of 0.

At each iteration, each patch is grown outwards, subject to the limitation that newly added cells must be within a given proximity to the cell. When multiple patches claim a cell, it is assigned to the patch with the highest value of proximity to the cell times patch “hunger”.

This algorithm does not converge at an optimum; rather, it quickly improves on the initial per-patch SSE(X), then wanders a space of comparable SSE(X) profiles until its final iteration.

Stratifying differential expression analyses across dozens or hundreds of patches leads to a huge number of models. To obtain a reasonable computation time, we ran each patch’s differential expression models using ordinary least squares rather than the more optimal smiDE. This approach abandons smiDE’s ability to adjust for bias from cell segmentation errors; however, this concern is ameliorated by the small size of each patch, which limits opportunities for confounding of this nature.

The PatchDE R code is available at https://github.com/Nanostring-Biostats/CosMx-Analysis-Scratch-Space/blob/Main/_code/PatchDE/DEutils.R.

---

**Supplemental Figure 1.**
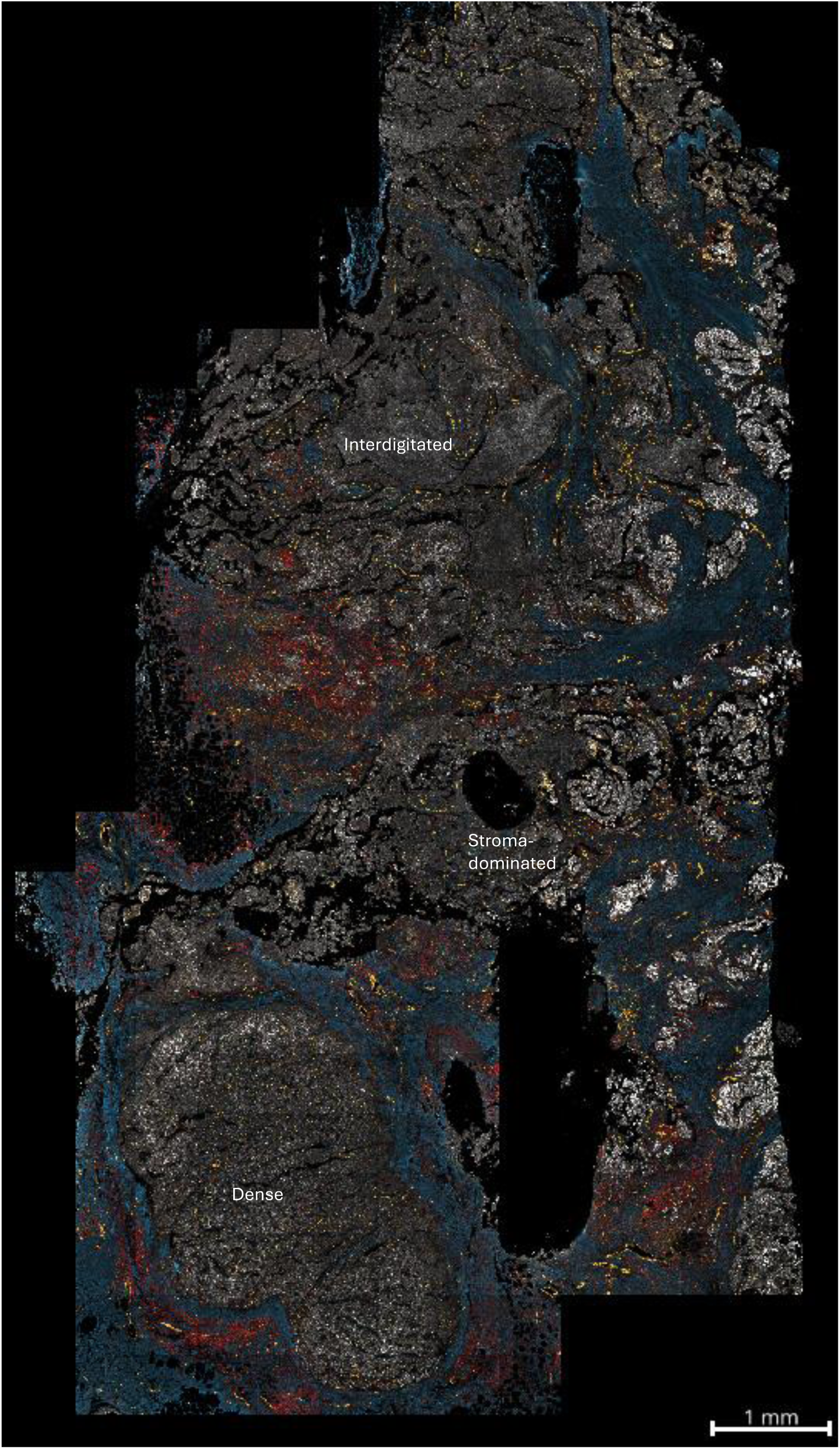
: protein landscape of a non-adjacent serial section of the same tumor. White: PanCK. Red: CD3. Blue: fibronectin. Yellow: CD31.

**Supplemental Figure 2:**
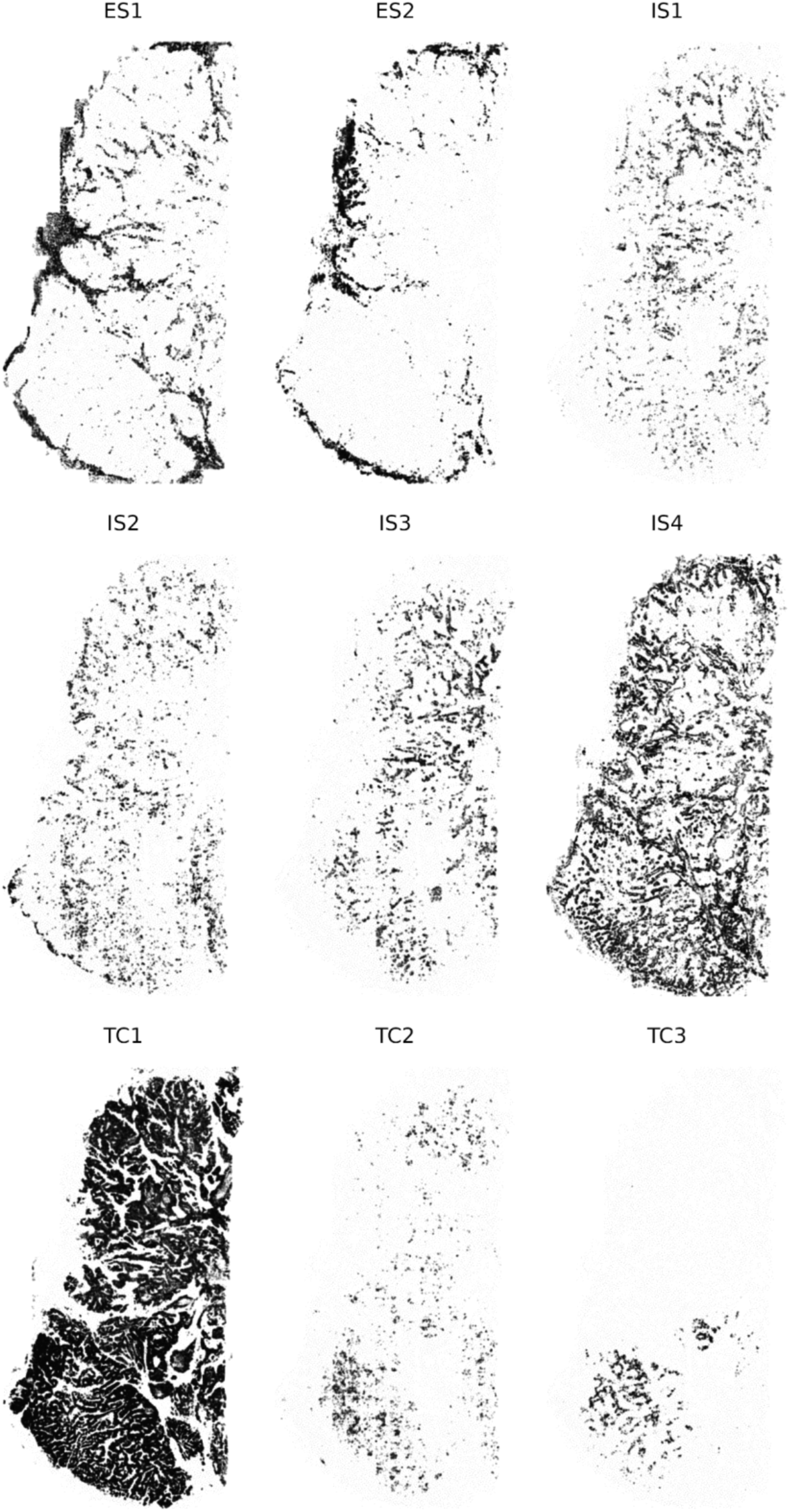
novae domains. Cells are colored by membership in each panel’s domain.

**Supplemental Figure 3:**
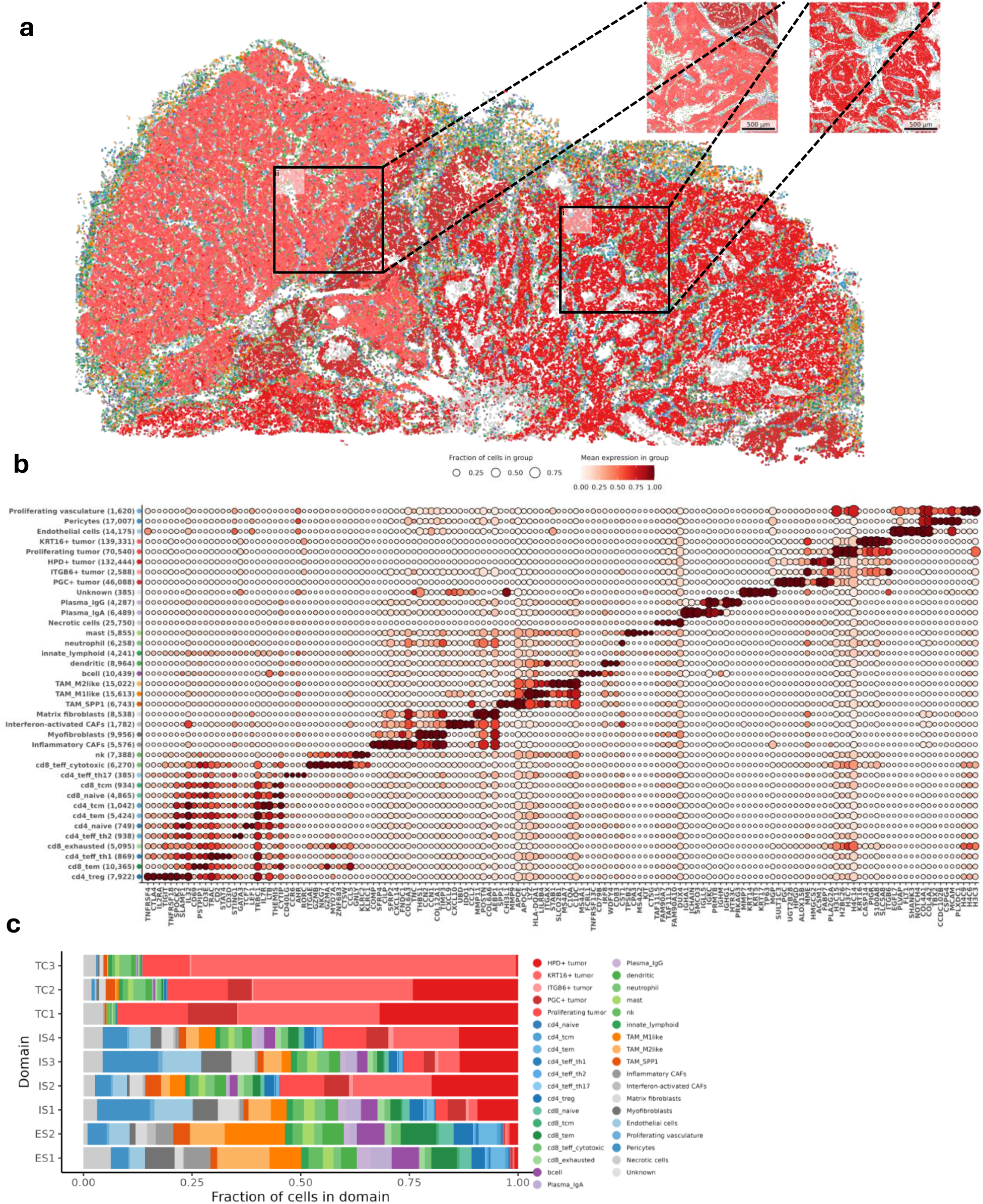
Cells divided into 37 minor cell types. a. Spatial plot of all minor cell types. Colors match those in (c). b. Marker heatmap shows top genes per cell type as well as total cells per cell type. c. Fraction of cells of each type found in each novae domain.

**Supplemental Figure 4:**
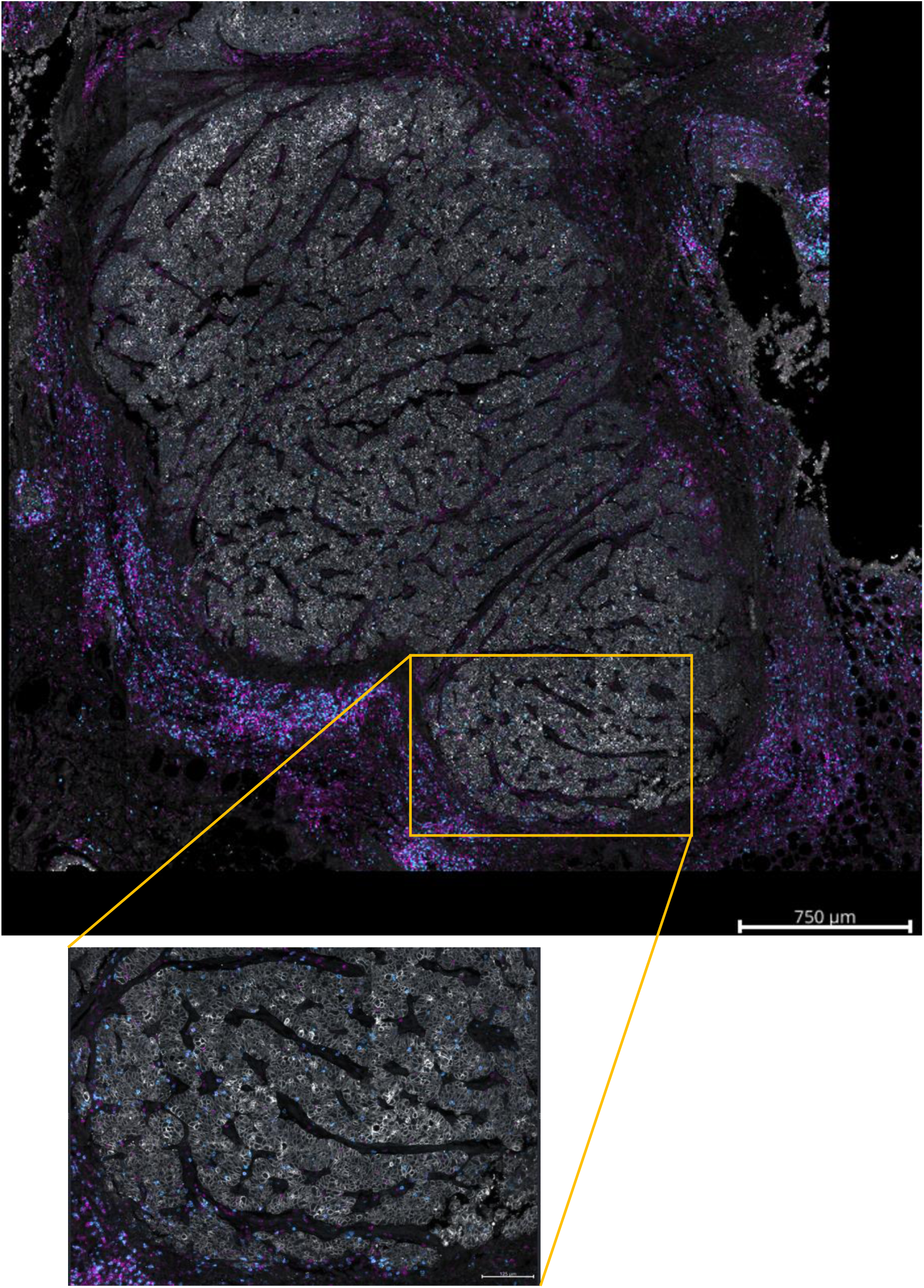
landscape of T-cell infiltrate around dense lobe. White: PanCK. Blue: CD8. Magenta: CD3.

**Supplemental Figure 5:**
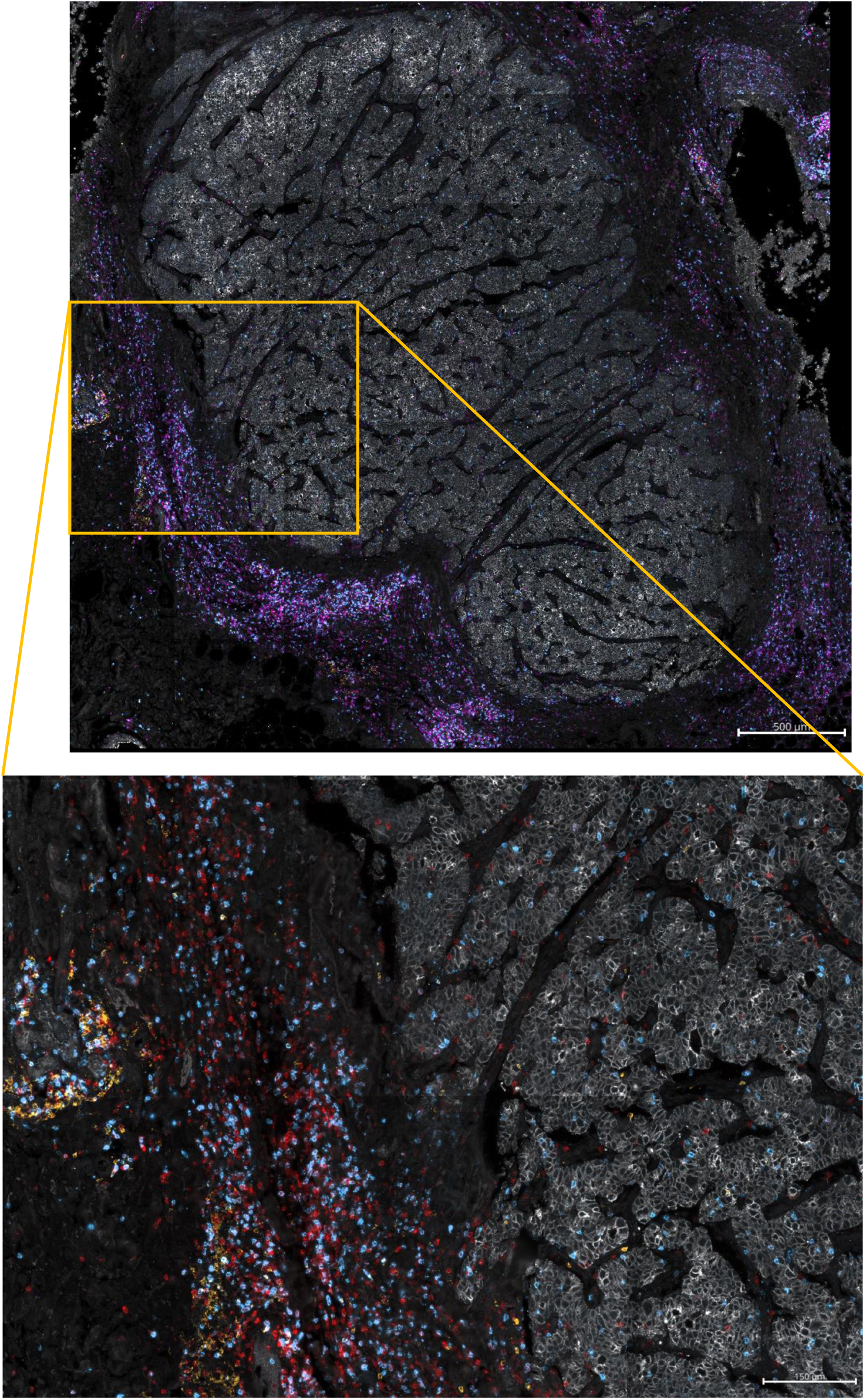
landscape of T-cell activation at right edge of dense lobe. White: PanCK. Red: CD3. Blue: CD8. Orange: CD45RA.

**Supplemental Figure 6:**
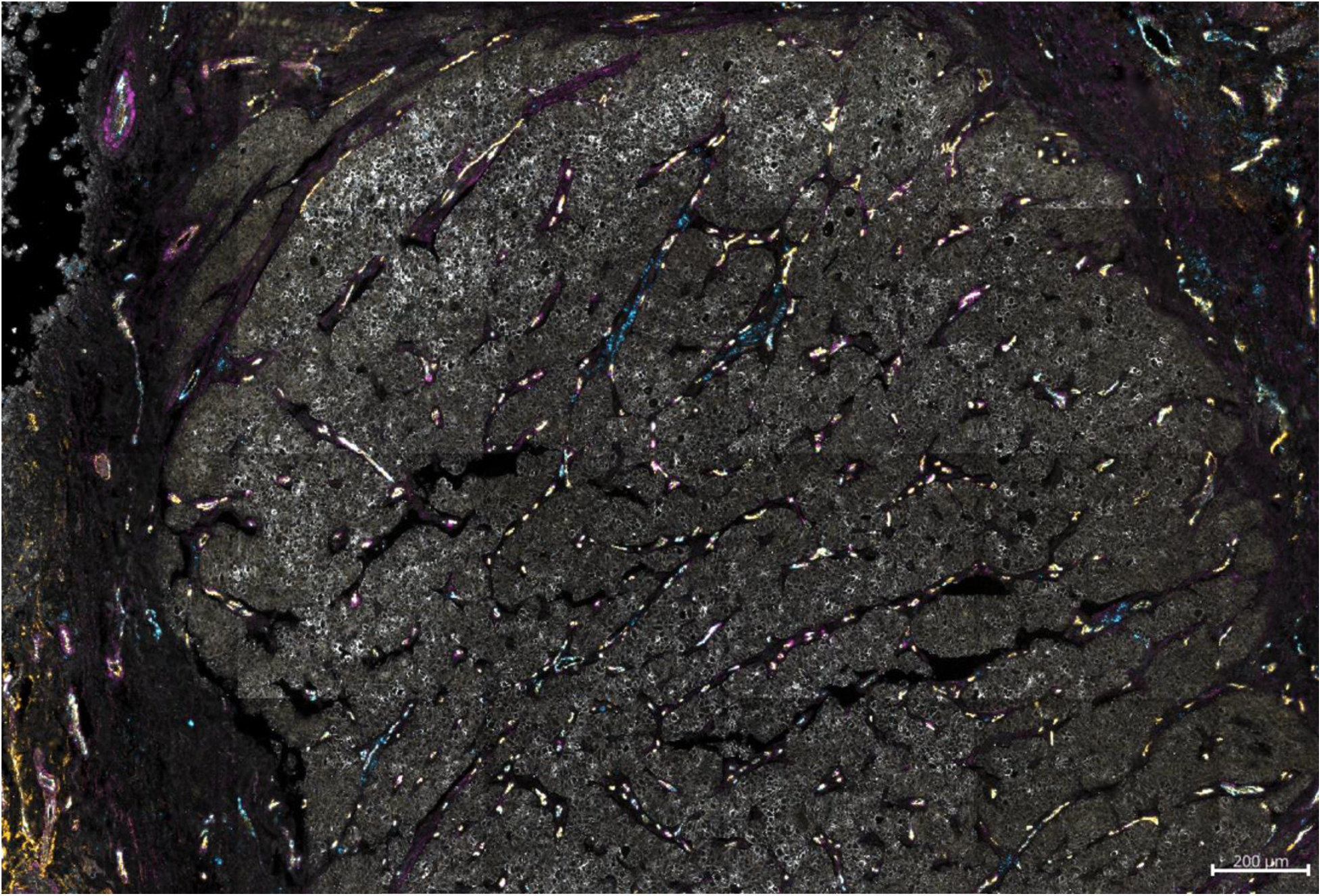
landscape of vasculature. White: PanCK. Purple: SMA. Blue: CD31. Yellow: CD34.

**Supplemental Figure 7:**
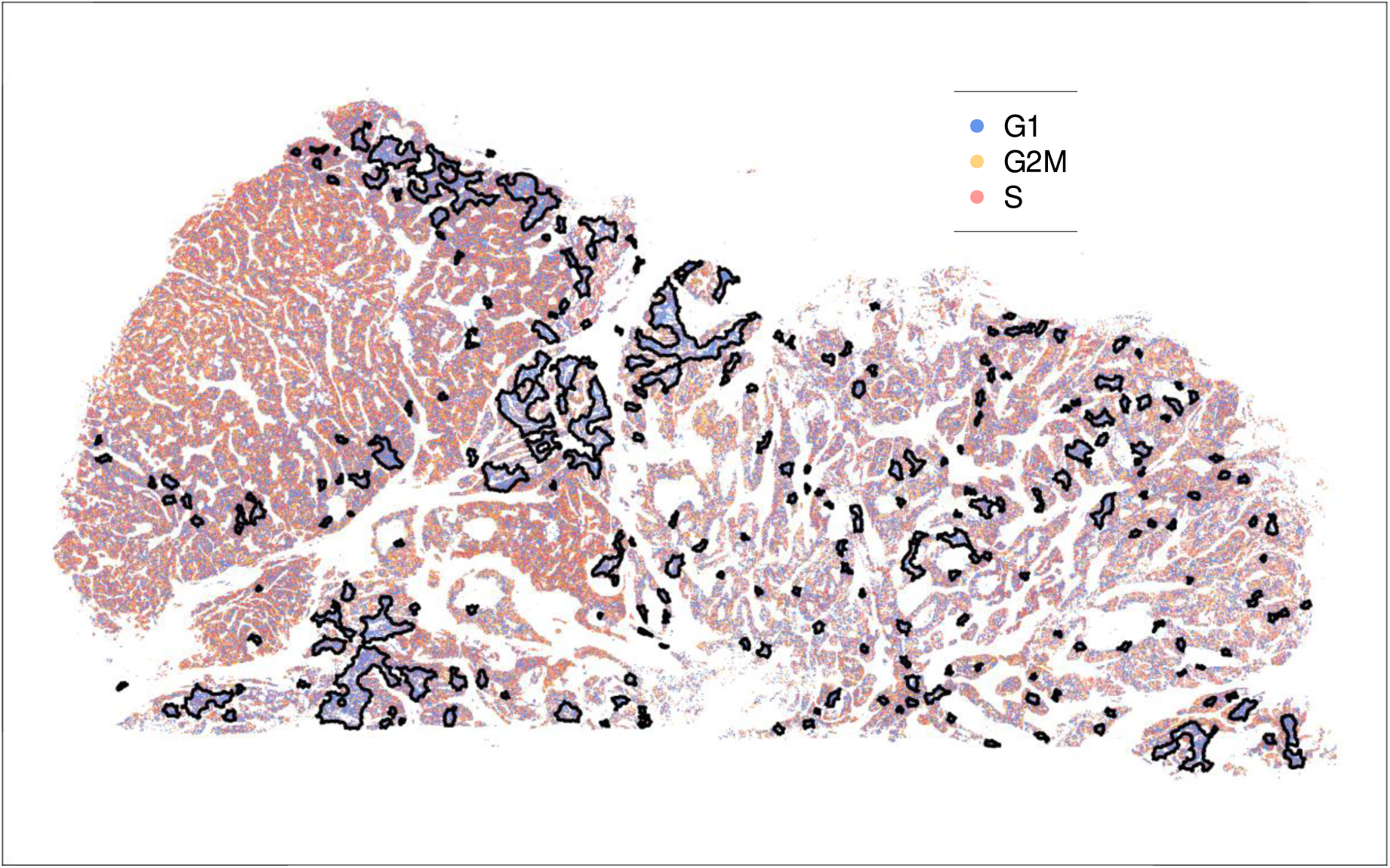
low-proliferation regions. Only tumor cells are shown. Color denotes cell cycle phase. Regions designated as lowproliferation are circled in black.

**Supplemental Figure 8:**
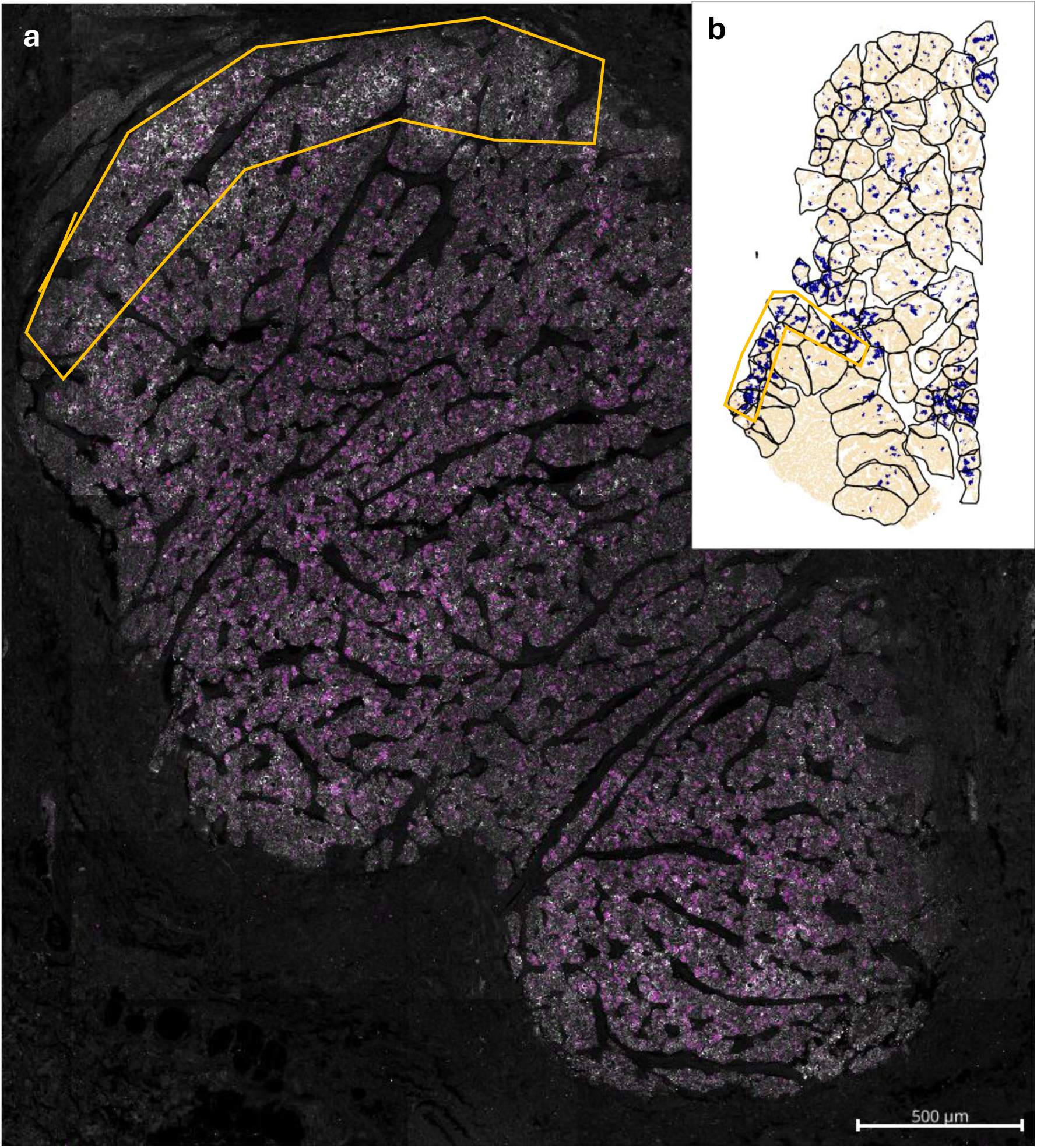
tumor proliferation. A. protein data. White: PanCK. Purple: Ki67. b. Low proliferation zones. From Fig 5, blue shows low proliferation regions. “Low proliferation edge” of dense lobe circled in both panels.

## References

1. Hanahan, D. & Weinberg, R. A. Hallmarks of Cancer: The Next Generation. Cell 144, 646–674 (2011).

2. Nanostring-Biostats. HieraType. https://github.com/Nanostring-Biostats/CosMx-Analysis-Scratch-Space/tree/Main/_code/HieraType (2025).

3. Wang, D. et al. Clinical Significance of Elevated S100A8 Expression in Breast Cancer Patients. Front. Oncol. 8, 496 (2018).

4. Elazezy, M. et al. Emerging Insights into Keratin 16 Expression during Metastatic Progression of Breast Cancer. Cancers 13, 3869 (2021).

5. Chen, J. et al. ALOX15B-based efferocytosis clusters: prognostic implications and immune cell infiltration in breast cancer. *Discov*. Oncol. 16, 1159 (2025).

6. Marino, N. et al. FAM83A is a potential biomarker for breast cancer initiation. Biomark. Res. 10, 8 (2022).

7. Feng, X. et al. CRABP2 regulates invasion and metastasis of breast cancer through hippo pathway dependent on ER status. J. Exp. Clin. Cancer Res. 38, 361 (2019).

8. Yu, W. et al. Increased expression of CYP4Z1 promotes tumor angiogenesis and growth in human breast cancer. Toxicol. Appl. Pharmacol. 264, 73–83 (2012).

9. Jorgovanovic, D., Song, M., Wang, L. & Zhang, Y. Roles of IFN-γ in tumor progression and regression: a review. Biomark. Res. 8, 49 (2020).

10. Escárcega, R. O., Fuentes-Alexandro, S., García-Carrasco, M., Gatica, A. & Zamora, A. The Transcription Factor Nuclear Factor-kappa B and Cancer. Clin. Oncol. 19, 154–161 (2007).

11. Si, W. et al. SARDH in the 1-C metabolism sculpts the T-cell fate and serves as a potential cancer therapeutic target. Cell. Mol. Immunol. 22, 1363–1378 (2025).

12. Levring, T. B. et al. Tumor necrosis factor induces rapid down-regulation of TXNIP in human T cells. Sci. Rep. 9, 16725 (2019).

13. Limoges, M.-A., Cloutier, M., Nandi, M., Ilangumaran, S. & Ramanathan, S. The GIMAP Family Proteins: An Incomplete Puzzle. Front. Immunol. 12, 679739 (2021).

14. De La Rosa Rodriguez, M. A. & Kersten, S. Regulation of lipid droplet homeostasis by hypoxia inducible lipid droplet associated HILPDA. Biochim. Biophys. Acta BBA - Mol. Cell Biol. Lipids 1865, 158738 (2020).

15. Luk, I. Y., Reehorst, C. M. & Mariadason, J. M. ELF3, ELF5, EHF and SPDEF Transcription Factors in Tissue Homeostasis and Cancer. Molecules 23, 2191 (2018).

16. Kar, A. & Gutierrez-Hartmann, A. ESE-1/ELF3 mRNA expression associates with poor survival outcomes in HER2+ breast cancer patients and is critical for tumorigenesis in HER2+ breast cancer cells. Oncotarget 8, 69622–69640 (2017).

17. Six, D. A. & Dennis, E. A. The expanding superfamily of phospholipase A2 enzymes: classification and characterization. Biochim. Biophys. Acta BBA - Mol. Cell Biol. Lipids 1488, 1–19 (2000).

18. Brand, A. et al. LDHA-Associated Lactic Acid Production Blunts Tumor Immunosurveillance by T and NK Cells. Cell Metab. 24, 657–671 (2016).

19. Chang, C.-H. et al. Metabolic Competition in the Tumor Microenvironment Is a Driver of Cancer Progression. Cell 162, 1229–1241 (2015).

20. Kay, E. J., Koulouras, G. & Zanivan, S. Regulation of Extracellular Matrix Production in Activated Fibroblasts: Roles of Amino Acid Metabolism in Collagen Synthesis. Front. Oncol. 11, 719922 (2021).

21. Pérez-Tomás, R. & Pérez-Guillén, I. Lactate in the Tumor Microenvironment: An Essential Molecule in Cancer Progression and Treatment. Cancers 12, 3244 (2020).

22. Cocciolone, A. J. et al. Elastin, arterial mechanics, and cardiovascular disease. Am. J. Physiol.-Heart Circ. Physiol. 315, H189–H205 (2018).

23. Westergaard, U. B., Andersen, M. H., Heegaard, C. W., Fedosov, S. N. & Petersen, T. E. Tetranectin binds hepatocyte growth factor and tissue-type plasminogen activator. Eur. J. Biochem. 270, 1850–1854 (2003).

24. Drexler, H. C. A. et al. Vascular Endothelial Receptor Tyrosine Phosphatase: Identification of Novel Substrates Related to Junctions and a Ternary Complex with EPHB4 and TIE2*[S]. Mol. Cell. Proteomics 18, 2058–2077 (2019).

25. Wu, L. & Zhu, Y. The function and mechanisms of action of LOXL2 in cancer (Review). Int. J. Mol. Med. 36, 1200–1204 (2015).

26. Corona, A. & Blobe, G. C. The role of the extracellular matrix protein TGFBI in cancer. Cell. Signal. 84, 110028 (2021).

27. Arpino, V., Brock, M. & Gill, S. E. The role of TIMPs in regulation of extracellular matrix proteolysis. Matrix Biol. 44–46, 247–254 (2015).

28. Christofk, H. R. et al. The M2 splice isoform of pyruvate kinase is important for cancer metabolism and tumour growth. Nature 452, 230–233 (2008).

29. Hamada, M. et al. Urokinase-Type Plasminogen Activator Receptor (uPAR) in Inflammation and Disease: A Unique Inflammatory Pathway Activator. Biomedicines 12, 1167 (2024).

30. Mustafa, S., Koran, S. & AlOmair, L. Insights Into the Role of Matrix Metalloproteinases in Cancer and its Various Therapeutic Aspects: A Review. Front. Mol. Biosci. 9, 896099 (2022).

31. Bylund, J., Hidestrand, M., Ingelman-Sundberg, M. & Oliw, E. H. Identification of CYP4F8 in Human Seminal Vesicles as a Prominent 19-Hydroxylase of Prostaglandin Endoperoxides. J. Biol. Chem. 275, 21844–21849 (2000).

32. Vainio, P. et al. Arachidonic Acid Pathway Members PLA2G7, HPGD, EPHX2, and CYP4F8 Identified as Putative Novel Therapeutic Targets in Prostate Cancer. Am. J. Pathol. 178, 525–536 (2011).

33. Baniwal, S. K., Chimge, N.-O., Jordan, V. C., Tripathy, D. & Frenkel, B. Prolactin-Induced Protein (PIP) Regulates Proliferation of Luminal A Type Breast Cancer Cells in an Estrogen-Independent Manner. PLoS ONE 8, e62361 (2013).

34. Urbaniak, A. et al. Prolactin-induced protein (PIP) increases the sensitivity of breast cancer cells to drug-induced apoptosis. Sci. Rep. 13, 6574 (2023).

35. Burrell, R. A., McGranahan, N., Bartek, J. & Swanton, C. The causes and consequences of genetic heterogeneity in cancer evolution. Nature 501, 338–345 (2013).

36. Mayo Clinic. TVB-2640 and Trastuzumab With Paclitaxel or Endocrine Therapy for Treatment of HER2 Positive Metastatic Breast Cancer. https://www.clinicaltrials.gov/study/NCT03179904.

37. Khafizov, R. et al. Sub-cellular Imaging of the Entire Protein-Coding Human Transcriptome (18933-plex) on FFPE Tissue Using Spatial Molecular Imaging. Preprint at 10.1101/2024.11.27.625536 (2024).

38. Bruker Spatial Biology. CosMx(R) SMI Manual Slide Preparation for Protein Assays. (2025).

39. He, S. et al. High-plex imaging of RNA and proteins at subcellular resolution in fixed tissue by spatial molecular imaging. Nat. Biotechnol. 40, 1794–1806 (2022).

40. napari contributors. napari: a multi-dimensional image viewer for python. (2019).

41. Nanostring-Biostats. napari-cosmx. https://github.com/Nanostring-Biostats/CosMx-Analysis-Scratch-Space/blob/Main/assets/napari-cosmx%20releases/napari_CosMx-0.4.17.4-py3-none-any.whl (2025).

42. Blampey, Q. et al. Novae: a graph-based foundation model for spatial transcriptomics data. Nat. Methods 22, 2539–2550 (2025).

43. Jassal, B. et al. The reactome pathway knowledgebase. Nucleic Acids Res. gkz1031 (2019) doi:10.1093/nar/gkz1031.

44. Aibar, S. et al. SCENIC: single-cell regulatory network inference and clustering. Nat. Methods 14, 1083–1086 (2017).

45. Vasconcelos, A. G., McGuire, D., Simon, N., Danaher, P. & Shojaie, A. Differential expression analysis for spatially correlated data. In press at Genome Biol (2025).

46. Korotkevich, G. et al. Fast gene set enrichment analysis. Preprint at 10.1101/060012 (2016).

